# Proteome-level robustness and the role of a histone-like protein during acute heat shock in the hyperthermophilic archaeon *Pyrococcus furiosus*

**DOI:** 10.1101/2025.05.02.651969

**Authors:** Haruko Okabe, Masahiro C. Miura, Asako Sato, Shungo Adachi, Akio Kanai

**Affiliations:** Institute for Advanced Biosciences, Keio University, Tsuruoka 997-0017, Japan; Faculty of Environment and Information Studies, Keio University, Fujisawa 252-0882, Japan; Systems Biology Program, Graduate School of Media and Governance, Keio University, Fujisawa 252-0882, Japan; National Cancer Center Research Institute, Tokyo 104-0045, Japan

**Author notes:** Corresponding Author Akio Kanai, PhD Institute for Advanced Biosciences, Keio University Tsuruoka, Yamagata 997-0017, Japan. Haruko Okabe and Masahiro C. Miura contributed equally to this work. The order of authorship depends on the time spent on the project.

**Keywords:** Archaea, Heat shock, Proteome, Transcriptome, Histone

## Abstract

The hyperthermophilic archaeon *Pyrococcus furiosus* thrives in extreme temperatures and exhibits a complex response to heat shock. However, the regulatory dynamics of genetic information during heat shock remain poorly understood. In this study, we exposed *P. furiosus* (cultured at 90°C) to acute heat shock by boiling (101–102°C) and analyzed its transcriptomic and proteomic responses. The levels of 16*S* and 23*S* rRNAs and of total tRNA were decreased by approximately 50%, and pre-tRNA splicing was inhibited, indicating suppression of translation. By contrast, approximately 90% of the proteome remained stable, underscoring the robustness of existing proteins. However, the transcriptome exhibited widespread alterations with limited correlation to the proteome (correlation coefficient *r* = 0.32), except for a few key proteins. These proteins included PF1883 (small heat shock protein), PF1385 (uracil-DNA glycosylase), and PF1616 (inositol-1-phosphate synthase), which are involved in protein chaperoning, stress-related metabolite synthesis, and DNA repair, respectively. Additionally, PF0624, previously annotated as a hypothetical protein, was identified as a putative histone motif-containing protein. Experimental evidence suggests that PF0624 may contribute to chromatin formation via archaeal histones in *P. furiosus*. In summary, our findings reveal that *P. furiosus* responds to acute heat shock by maintaining protein stability, suppressing translation, limiting genomic damage, and potentially compacting genomic DNA into archaeal chromatin.

**IMPORTANCE:** Hyperthermophilic archaea, such as *Pyrococcus furiosus*, thrive in extreme environments where the temperatures may reach up to 100°C. However, the precise mechanisms by which these organisms protect their genomic DNA from heat-induced damage remain incompletely understood. In this study, we propose that PF0624, a histone-like protein that is transcriptionally induced and translated in response to acute heat shock, is critical in stabilizing archaeal chromatin structure through histone-mediated mechanisms. Our results highlight the sophisticated molecular strategies employed by *P. furiosus* to survive extreme thermal stress.

## INTRODUCTION

Microorganisms that inhabit extreme environments, particularly archaea, have long served as valuable models for studying biological adaptation and stress responses in harsh conditions. These extremophiles thrive in environments with temperatures exceeding 100°C (1), hypersaline conditions with NaCl concentrations of up to 5 M (2), or highly acidic surroundings with pH values as low as 1–2 (3). In such studies, extreme environmental parameters, such as high temperature, high salt concentration, oxidative stress, or radiation, are often examined as stress factors, and the corresponding molecular responses are analyzed to gain insight into the underlying adaptive mechanisms. The heat shock response in hyperthermophilic archaea, particularly the study of thermostable proteins, was a major focus of research in the 1990s and early 2000s. Much of this work centered on heat shock proteins, especially molecular chaperones, following the groundwork laid in bacterial systems (4–6). From the early 2000s onward, the field advanced toward identifying transcription factors, such as Phr, which commonly regulate heat-inducible genes (7, 8). This shift facilitated the transition to system-level investigations using omics-based approaches, including transcriptomics (global transcriptional profiling), proteomics (comprehensive protein analysis), and metabolomics (systematic profiling of metabolites), to achieve a holistic understanding of microbial physiology under stress conditions.

*Pyrococcus furiosus*, a hyperthermophilic archaeon, was first isolated in the 1980s from a sulfur-rich hydrothermal vent on the island of Vulcano, Italy (9). This archaeon grows at temperatures ranging from 70 to 103°C, with an optimal growth temperature of 100°C (10). In hydrothermal vent environments, although ambient temperatures can reach 100°C, seawater does not boil due to the high hydrostatic pressure at depth. Following the publication of its complete genome sequence in 2001 (11), *P. furiosus* became a common model organism for systems biology, particularly in studies using transcriptomic and proteomic analyses. For example, cDNA microarrays have been used to characterize gene expression under conditions of heat shock and oxidative stress (12, 13); proteomics has revealed insights into protein complexes (14); and integrated multi-omics approaches, combining transcriptomics, proteomics, and metabolomics, have expanded our understanding of the organism’s cellular dynamics (15, 16). Our group has also contributed to this field through systematic identification of RNA-associated enzymes, using expression library screening (17–19) and bioinformatics methods based on machine learning to predict the functions of all genome-encoded proteins (20).

In this study, we examined the cellular response of *P. furiosus* to acute heat shock, in which the culture medium was boiled at 101–102°C under atmospheric pressure. Proteomic and RNA sequencing (RNA-seq) analyses were performed on the same cell pellets, enabling direct comparison of transcriptional and translational responses. Under these conditions, the transcription profile exhibited substantial fluctuations, whereas translation was globally suppressed. Approximately 90% of the pre-existing proteins remained largely unchanged, with only a few exceptions. One such exception was the cell division protein FtsZ2, which was rapidly degraded, suggesting that cell division is inhibited under acute heat shock. By contrast, the *PF0624* gene, which encodes a histone-fold protein, was upregulated at the transcript and protein levels, implying that the protein plays a role in promoting chromatin organization during heat shock in *P. furiosus*.

## RESULTS AND DISCUSSION

### Heat shock leads to a reduction in total tRNA levels and inhibition of pre-tRNA splicing in the hyperthermophilic archaeon *P. furiosus*

*P. furiosus* cultures were maintained at 90°C as a control condition, and heat shock conditions were induced by incubating the cultures at boiling temperature (corresponding to 101–102°C at 1 atm) for 15, 30, and 45 min. Under these conditions, we observed a time-dependent decrease in the total tRNA levels, with reduction of approximately 50% after 45 min of boiling (Fig. 1A). To investigate this in more detail, we selected tRNA^Trp^(CCA) as a representative tRNA and analyzed its expression by northern blotting. Consistent with the global tRNA trend, the level of tRNA^Trp^(CCA) decreased to about half of its original level after 45 min of heat shock (Fig. 1B). The *P. furiosus* genome contains two intron-containing tRNA genes, *tRNA^Trp^(CCA)* and *tRNA^Met^(CAU)* (Fig. 1C). To assess the effect of heat shock on pre-tRNA splicing, we conducted reverse transcription (RT)-PCR analysis. The results showed heat-dependent accumulation of pre-tRNA species for both genes (Fig. 1D). Quantitative PCR analysis revealed that the levels of pre-tRNA^Trp^(CCA) and pre-tRNA^Met^(CAU) increased by approximately 12- and 17-fold, respectively, under heat shock conditions (Fig. 1E). The mature tRNAs could not be detected by RT-PCR with the primers used in this study, regardless of the culture temperature. This is likely due to the presence of chemical modifications in tRNAs that interfere with the RT reaction (21). Taken together, these results suggest that heat shock inhibits pre-tRNA splicing and increases the degradation of mature tRNAs, thereby decreasing total tRNA levels in *P. furiosus*. Consequently, these conditions are likely to impair mRNA translation as a potential mechanism by which heat shock affects gene expression at the translational level.

**FIG 1.**
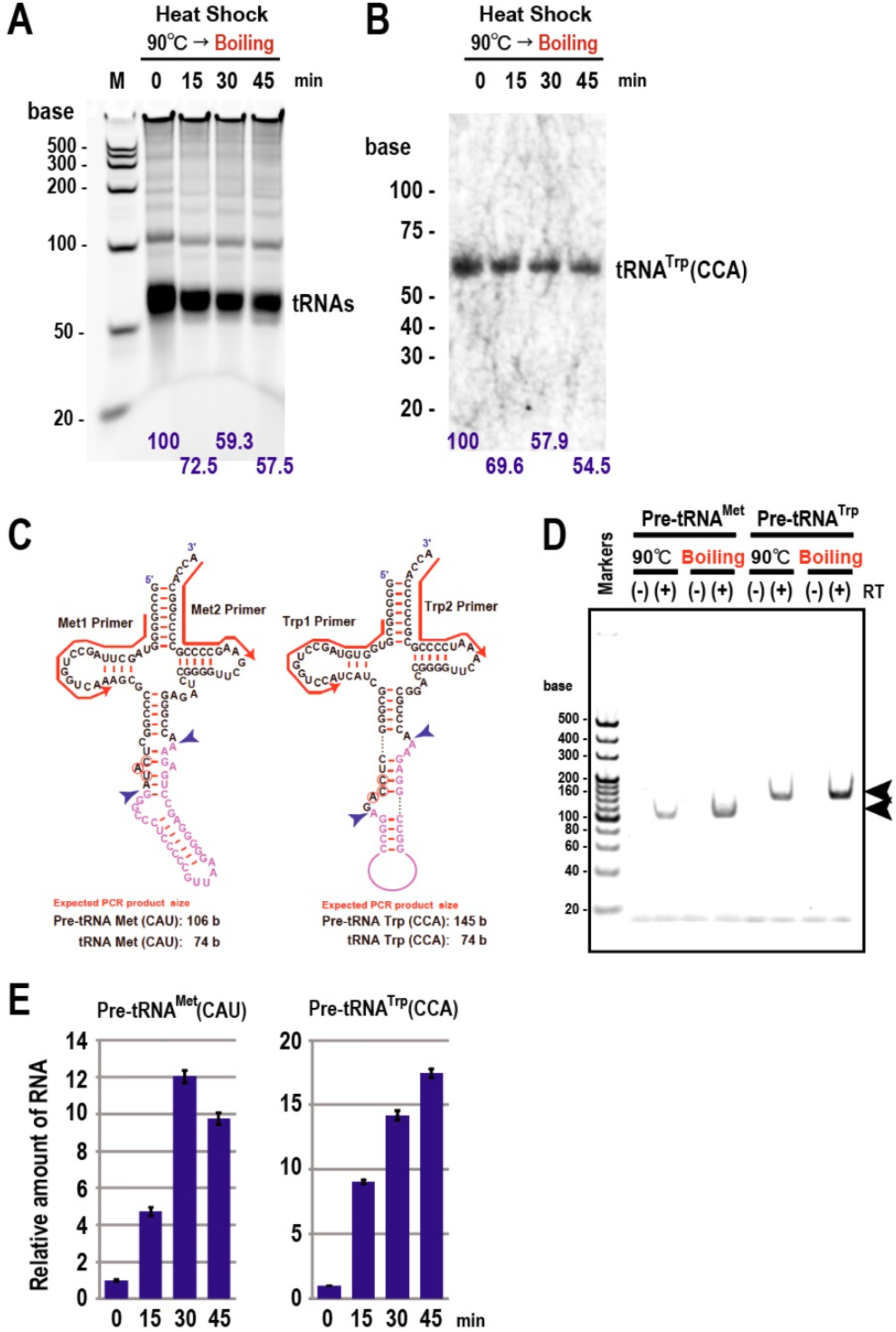
Decrease in tRNA levels and inactivation of pre-tRNA splicing in *P. furiosus* exposed to heat shock conditions. (A) Decrease in *P. furiosus* tRNA levels following heat shock. RNA (4 μg) from each timepoint (0 min (90°C), and 15, 30, and 45 min after boiling (101–102°C) began) were subjected to denaturing gel electrophoresis on a 1% agarose gel and stained with SYBR Green II. The relative amounts of tRNA are indicated at the bottom of each lane. Similar results were obtained in at least three independent experiments. (B) Northern blotting of tRNA^Trp^(CCA) after heat shock. The relative amounts of tRNA^Trp^(CCA) are indicated at the bottom of each lane. Pre-tRNA^Trp^(CCA) was not detected under these conditions (see text for details). (C) Sequences and RNA secondary structures of pre-tRNA^Met^(CAU) and pre-tRNA^Trp^(CCA). Exons are shown in black and introns in pink. The anticodon sequences are circled in red. Blue arrowheads indicate exon–intron boundaries cleaved by splicing endonuclease. Red arrows indicate primer positions for RT-PCR analysis. (D) Accumulation of pre-tRNA^Met^(CAU) and pre-tRNA^Trp^(CCA) in *P. furiosus* exposed to heat shock conditions. RT-PCR analysis of pre-tRNA^Met^(CAU) and pre-tRNA^Trp^(CCA) was performed using total RNA extracted from *P. furiosus* incubated either at 90°C or after boiling for 15 min. Arrowheads indicate the positions of each pre-tRNA. (+)/(−) RT: reactions with and without reverse transcriptase. Mature tRNAs were not amplified under these conditions (see text for details). (E) RT-qPCR analysis of pre-tRNA^Met^(CAU) and pre-tRNA^Trp^(CCA) under heat shock conditions. RT-qPCR analysis of the indicated RNAs was performed using total RNA extracted from *P. furiosus* at each timepoint. The mean (*n* = 3) and standard deviation are shown.

### Despite heat shock, the proteome of *P. furiosus* remains largely stable, whereas the transcriptome is substantially disrupted

Given the inhibition of the translation process in *P. furiosus* under heat shock conditions, we next investigated changes at the proteomic level under the four conditions described in the previous section (Fig. 2). First, sodium dodecyl sulfate-polyacrylamide gel electrophoresis (SDS-PAGE) analysis of the total protein extracts revealed no notable changes in the overall protein levels following heat shock (Fig. 2A). This observation was supported by quantitative proteome analysis, which showed that 94.5% of the 925 identified proteins after 15 min of heat shock, 92.6% after 30 min, and 88.3% after 45 min did not exhibit significant changes (defined as an increase by at least twofold or a decrease to less than half of the control level) compared with the control sample (Fig. 2B‒D). These findings indicate that the *P. furiosus* proteome is remarkably robust against thermal stress. Nonetheless, some proteins did show marked changes in their abundance. Four proteins were significantly upregulated, with increases of approximately three-to 12-fold after 45 min of heat shock compared with the control. Conversely, two proteins were drastically downregulated, with levels reduced to approximately 1/44 to 1/20 after 30 min of heat shock. According to annotations in the RefSeq *P. furiosus* DSM 3638 protein database (National Center for Biotechnology Information [NCBI]), the upregulated proteins were identified as PF0624 (hypothetical protein), PF1385 (type-4 uracil-DNA glycosylase), PF1616 (*myo*-inositol-1-phosphate synthase), and PF1883 (small heat shock protein). The downregulated proteins were PF0525 (cell division protein FtsZ2) and PF0798 (glycosyl transferase). Because the majority of proteins did not respond significantly to heat shock, no substantial shifts were observed in the Archaeal Clusters of Orthologous Genes (arCOG) functional classifications (Supplementary Table S1).

**FIG 2.**
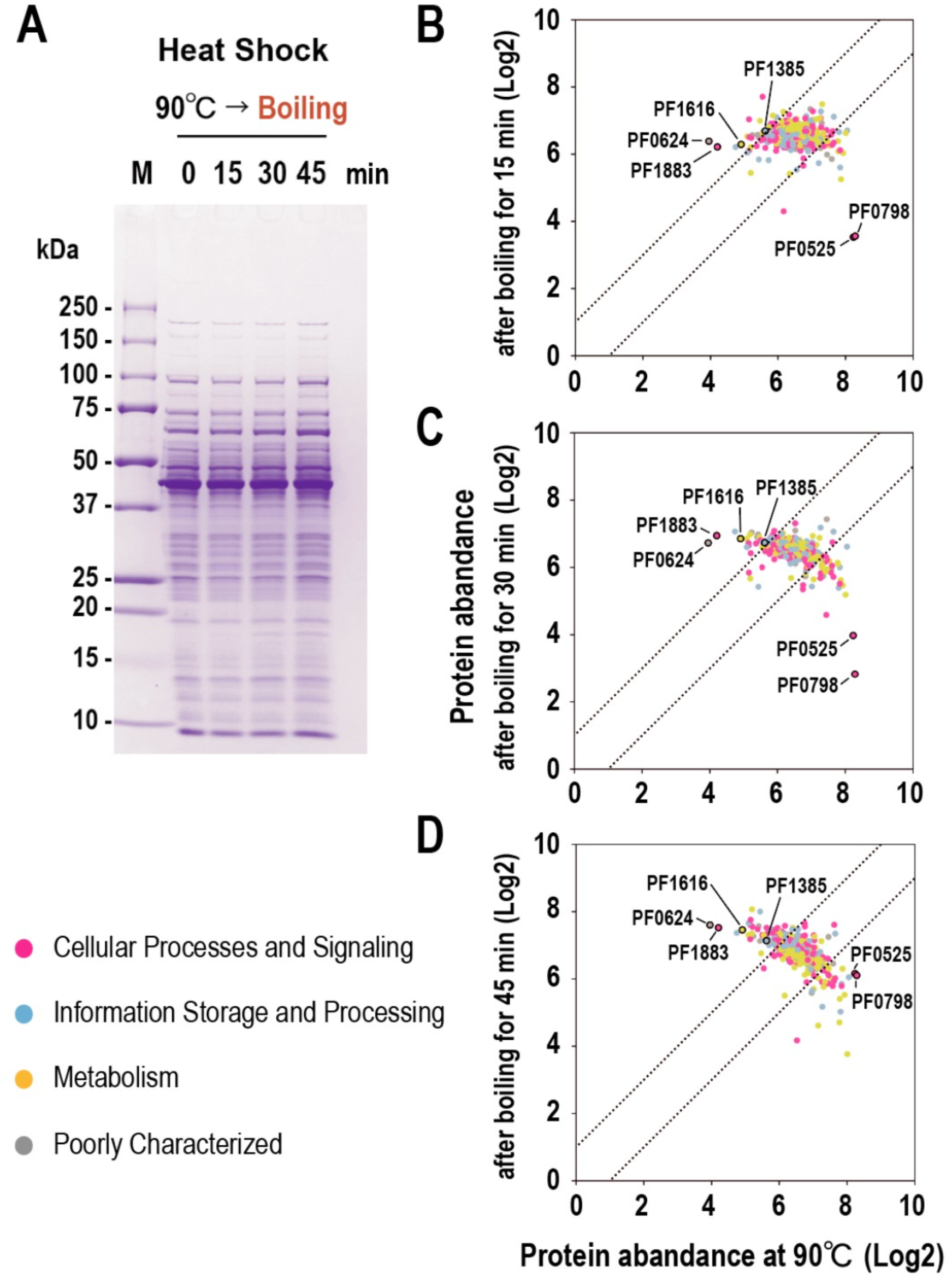
*P. furiosus* proteomic profile remains largely unchanged after heat shock. (A) SDS-PAGE (10%–20% polyacrylamide gels) of the total extracted proteins from *P. furiosus* before (90°C) and after boiling the culture medium (101–102°C). Proteins were stained with Coomassie brilliant blue. (B–D) Comparison of the levels of 927 proteins before and after heat shock for 15 (B), 30 (C), and 45 min (D). Each dot represents an individual protein, colored according to the arCOG general functional classification (see Supplementary Table S1). The dashed lines indicate the threshold for a twofold change. The positions of six notable proteins are labeled: PF0525, cell division protein FtsZ2; PF0624, hypothetical protein; PF0798, glycosyltransferase family 2 protein; PF1385, type-4 uracil-DNA glycosylase; PF1616, *myo*-inositol-1-phosphate synthase; and PF1883, small heat shock protein.

In contrast to the relatively stable proteomic profile, the transcriptomic landscape of *P. furiosus* displayed substantial changes in response to heat shock (Fig. 3). Denaturing gel electrophoresis of total RNA revealed that, after 15 min of heat exposure, the levels of 23*S* and 16*S* rRNAs were reduced to approximately 40% and 60% of their control levels, respectively (Fig. 3A,B). This pattern closely resembled the heat shock-induced decrease in tRNA levels (Fig. 1), suggesting a common mechanism affecting multiple classes of structural RNAs during thermal stress. At the transcriptome level, out of 1983 detectable transcripts, 1138 (57.4%) were upregulated by more than twofold after 15 min of heat shock. This number increased to 1237 transcripts (62.4%) after 30 min and remained high at 1217 transcripts (61.4%) after 45 min (Fig. 3C‒E). The number of transcripts that remained unchanged across these conditions averaged 651 (32.8%), and only about 135 transcripts (6.6%) increased by more than twofold. Many of the downregulated transcripts were associated with metabolic functions, based on the arCOG functional classification (Supplementary Table S2). For example, among the 130 transcripts that decreased by more than half after 15 min of heat shock, 78 (60%) were classified as metabolism-related genes. Of these, 43 belonged to category (C) “Energy production and conversion,” indicating that genes involved in core metabolic pathways are particularly susceptible to transcriptional repression under heat shock conditions. These findings highlight a marked contrast between the heat-induced stability of the proteome and the dynamic, stress-responsive nature of the *P. furiosus* transcriptome. The rapid downregulation of rRNA and metabolic genes suggests a coordinated transcriptional response aimed at conserving energy and reallocating resources under extreme environmental stress.

**FIG 3.**
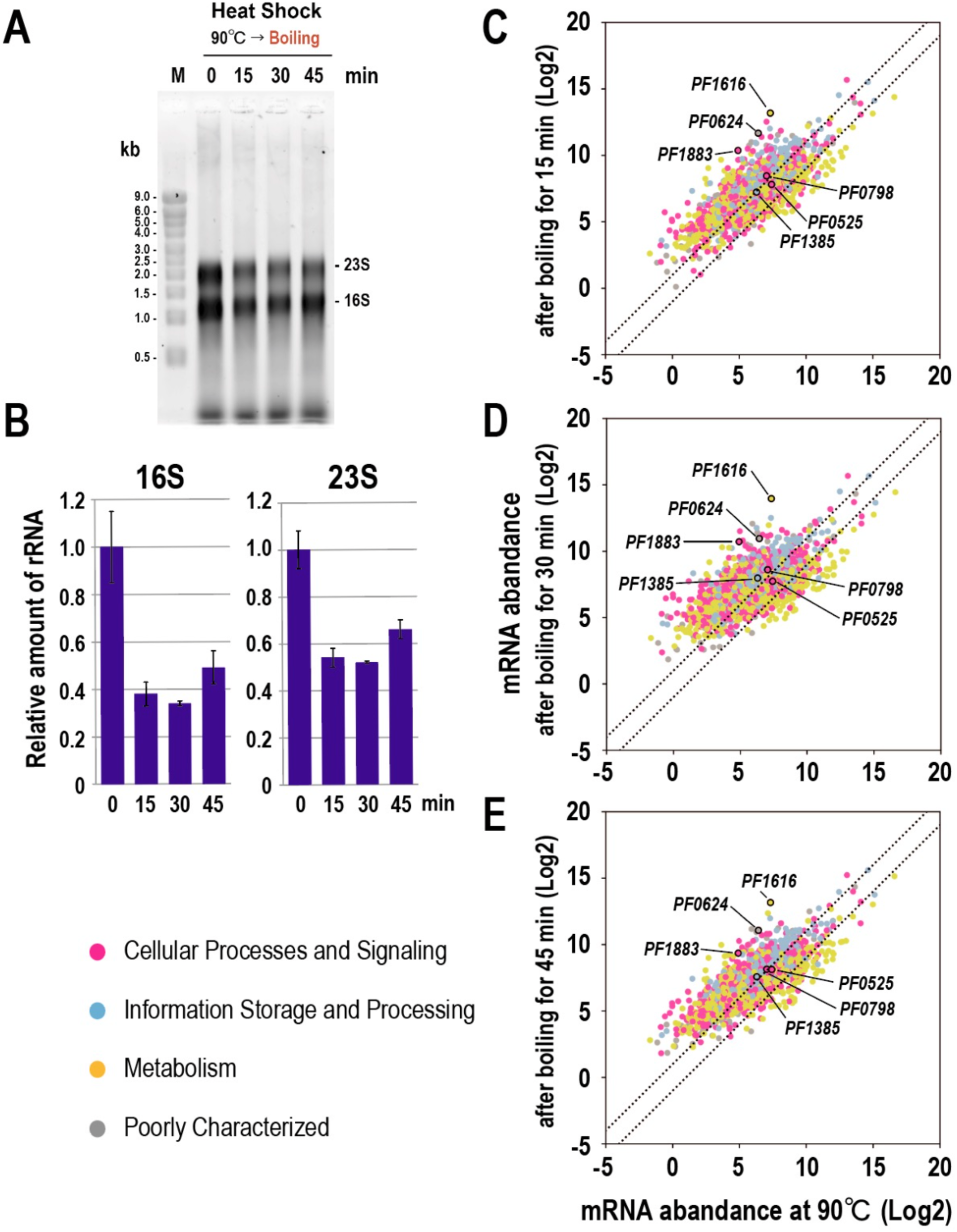
*P. furiosus* transcriptome is significantly disrupted by heat shock. (A) Denaturing gel electrophoresis of the total RNAs from *P. furiosus* used for RNA-seq. Total RNA prepared from *P. furiosus* before (90°C) and after boiling the culture medium (101‒102°C) were subjected to denaturing gel electrophoresis on a 1% agarose gel and stained with SYBR Green II. (B) qPCR analysis of 16*S* and 23*S* rRNAs under heat shock conditions. qPCR analysis of the indicated RNAs was performed using total RNA extracted from *P. furiosus* at each timepoint. The mean (*n* = 3) and standard deviation are shown. (C‒E) Comparison of the levels of 2070 mRNAs before and after heat shock for 15 (C), 30 (D), and 45 min (E). Each dot corresponds to an individual mRNA, colored according to the arCOG general functional classification (see Supplementary Table S2). The dashed lines delineate the threshold for a twofold change. The positions of six notable mRNAs are indicated (see Fig. 2 for protein annotation).

An analysis of 925 genes detected in both the transcriptomic and proteomic datasets revealed a poor correlation between the changes in transcript and protein levels, with only a few exceptions (*r* = 0.32 at 15 min post heat shock; Supplementary Fig. S1). This indicates that, although heat shock broadly induces mRNA expression, the translation of most transcripts does not occur, which is likely due to the impairment of the translation machinery. Among the few protein genes whose expression levels were correlated with mRNA induction were *PF0624*, *PF1385*, *PF1616*, and *PF1883* (Supplementary Fig. S1 and S2). Excluding *PF0624*, which encodes a hypothetical protein, the upregulation of the remaining three proteins can be explained based on their known biological functions (Supplementary Fig. S2). *PF1883* encodes a small heat shock protein, a chaperone induced by heat shock that prevents aggregation and aids the refolding of thermally denatured proteins (22). *PF1385* encodes a type-4 uracil-DNA glycosylase, an enzyme involved in DNA repair that likely excises uracil residues formed by cytosine deamination during heat shock (23, 24). *PF1616* encodes *myo*-inositol-1-phosphate synthase, which synthesizes di-*myo*-inositol-1-phosphate, a compound that protects cells from thermal damage (25, 26). Except for *PF1385*, the genes *PF0624*, *PF1616*, and *PF1883* are transcriptionally activated downstream of the heat-responsive transcription factor Phr (*PF1790*), whose repression is relieved by heat shock (7, 8). Consistent with this, the transcript levels of these three genes increased by approximately 40‒100-fold after heat shock. As previously described, *PF0525* (cell division protein FtsZ2) and *PF0798* (glycosyl transferase) were dramatically reduced at the protein level following heat shock. Given the role of FtsZ2 in cell division, this reduction implies that heat shock suppresses the cell division process in *P. furiosus* (27). According to STRING database annotations (28), *PF0798* encodes a glycosyl transferase associated with cell envelope functions (Function Code: 3.2), including biosynthesis of surface polysaccharide, lipopolysaccharide, and antigens. Thus, this enzyme may also play a role in cell division through membrane-associated processes, although further experimental validation is required. Interestingly, the mRNA levels of *PF0525* and *PF0798* remained relatively unchanged before and after heat shock, whereas their corresponding protein levels were significantly reduced (Fig. 2 and 3, and Supplementary Fig. S2). This discrepancy suggests the presence of heat shock-triggered, protein-specific degradation mechanisms that may be independent of transcriptional regulation.

### Structure and putative function of the hypothetical protein PF0624

To determine the function of PF0624, we performed structural modeling of the protein because its relatively short length (153 amino acids) made it suitable for such analysis. Using Phyre2 (29) for structure prediction, PF0624 was inferred to possess a histone-fold motif (Fig. 4A). A search for structurally similar proteins in the Protein Data Bank (PDB) revealed that the three-dimensional structure of archaeal histone B from *Methanothermus fervidus* (69 amino acids) was previously resolved (30) (Fig. 4B, shown as a dimer). This protein shares high sequence conservation with archaeal histone B from *P. furiosus* (67 amino acids), with 57% identity and 92% similarity at the amino acid level. Comparison of the predicted structure of PF0624 (monomeric form) and the known dimeric structure of archaeal histone B revealed a striking structural similarity (Fig. 4C). This finding suggests that a single polypeptide of PF0624 adopts a structure closely resembling that of a histone dimer, indicating that PF0624 may function in a histone-like manner despite its monomeric architecture. Recent computational studies on archaeal histone-like proteins (31) have classified PF0624-like proteins with histone-fold motifs into the “bacteria-type doublets” group, which appears to be distributed among members of the hyperthermophilic order *Thermococcales*. Furthermore, in *Methanocaldococcus jannaschii*, the histone variant MJ1647 forms a tetramer and bind to DNA (32). However, MJ1647 is only 96 amino acids long, considerably shorter than PF0624 (153 amino acids). Amino acid sequence alignment between PF0624 and MJ1647 revealed limited sequence identity (17.7%; 17/96 residues) but moderate similarity (57.2%; 55/96 residues), indicating potential functional divergence despite shared structural features.

**FIG 4.**
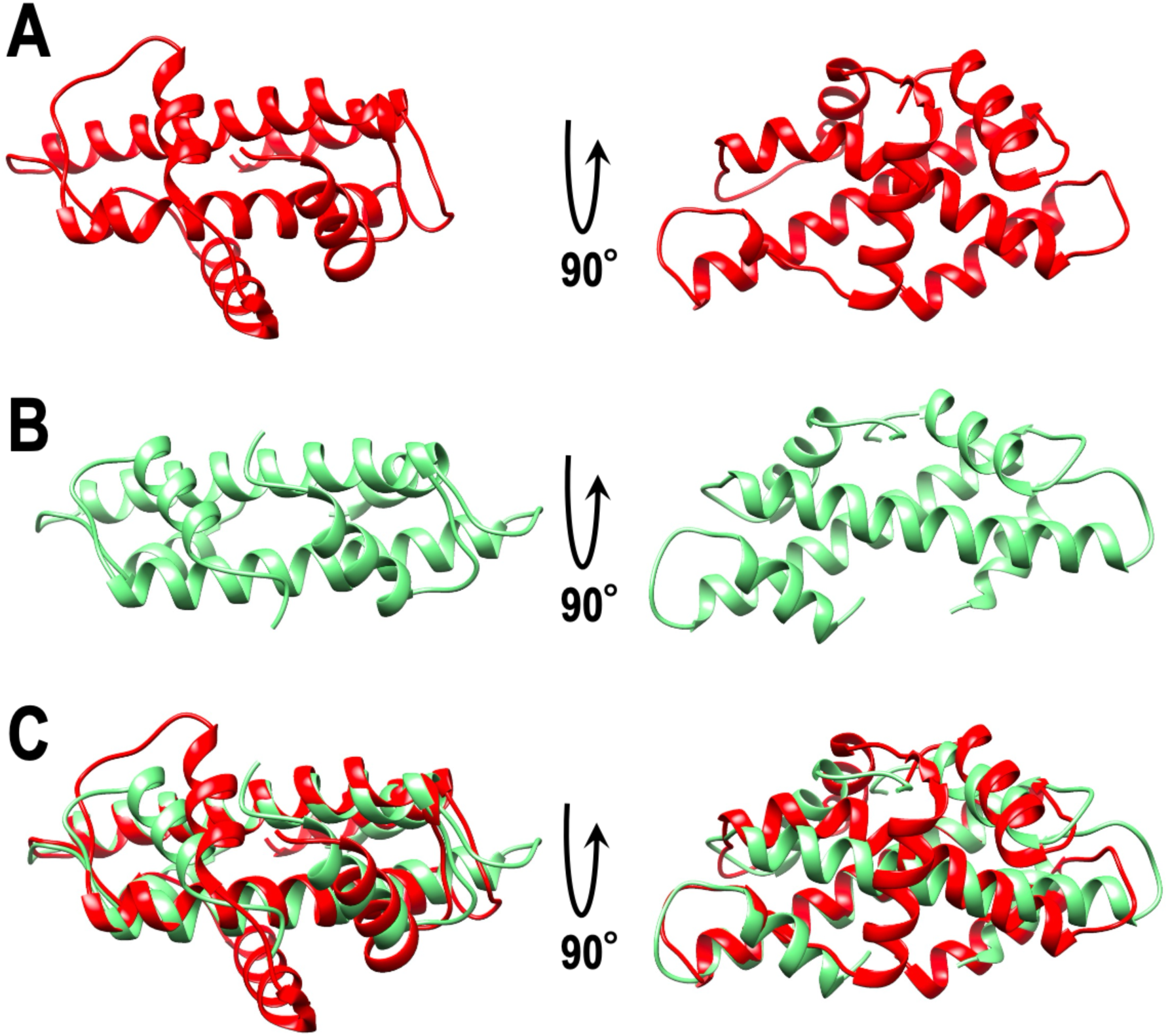
Comparison of the tertiary structures of PF0624 protein (histone-fold protein) and an archaeal histone. (A) Three-dimensional views of predicted structure of PF0624 protein (monomer structure). (B) Three-dimensional structure of archaeal histone B (dimer structure) from *M. fervidus* (PDB ID: 5T5K). (C) Merged image.

To investigate whether PF0624 possesses DNA-binding activity similar to that of archaeal histones, we prepared recombinant forms of both PF0624 and *P. furiosus* histone (*Pf*-histone) without affinity tags, such as His-tags, to eliminate any interference with the native binding properties of the proteins. For *Pf*-histone, we initially added a His-tag to the N-terminus to facilitate purification via Ni²⁺ affinity chromatography, followed by cleavage of the tag using factor Xa protease to yield the native form (Fig. 5A,B). By contrast, PF0624 was expressed at high levels in *E. coli*, allowing us to purify the recombinant protein with no affinity tags. After harvesting total protein, heat treatment was followed by anion exchange chromatography, which yielded a high-purity protein, as judged by SDS-PAGE (Fig. 5C). Using these purified tag-free proteins, we evaluated their ability to interact with DNA using a biosensor system, which quantitatively measures biomolecular interactions based on nanogram-level frequency shifts of a quartz resonator. At 45°C (the upper temperature limit for the biosensor system is 50°C), PF0624 exhibited clear binding to double-stranded DNA (dsDNA), comparable to that observed for the positive control, *Pf*-histone (Fig. 5D). As expected, bovine serum albumin (BSA), used as a negative control, showed negligible interaction. PF0624 also interacted directly with *Pf*-histone (Fig. 5E), whereas no significant interaction was observed between *Pf*-histone and BSA. These results suggest that PF0624 not only binds to DNA, like archaeal histones, but may also participate in histone-like protein complexes, highlighting its potential role in chromatin organization or DNA-related processes in *P. furiosus*.

**FIG 5.**
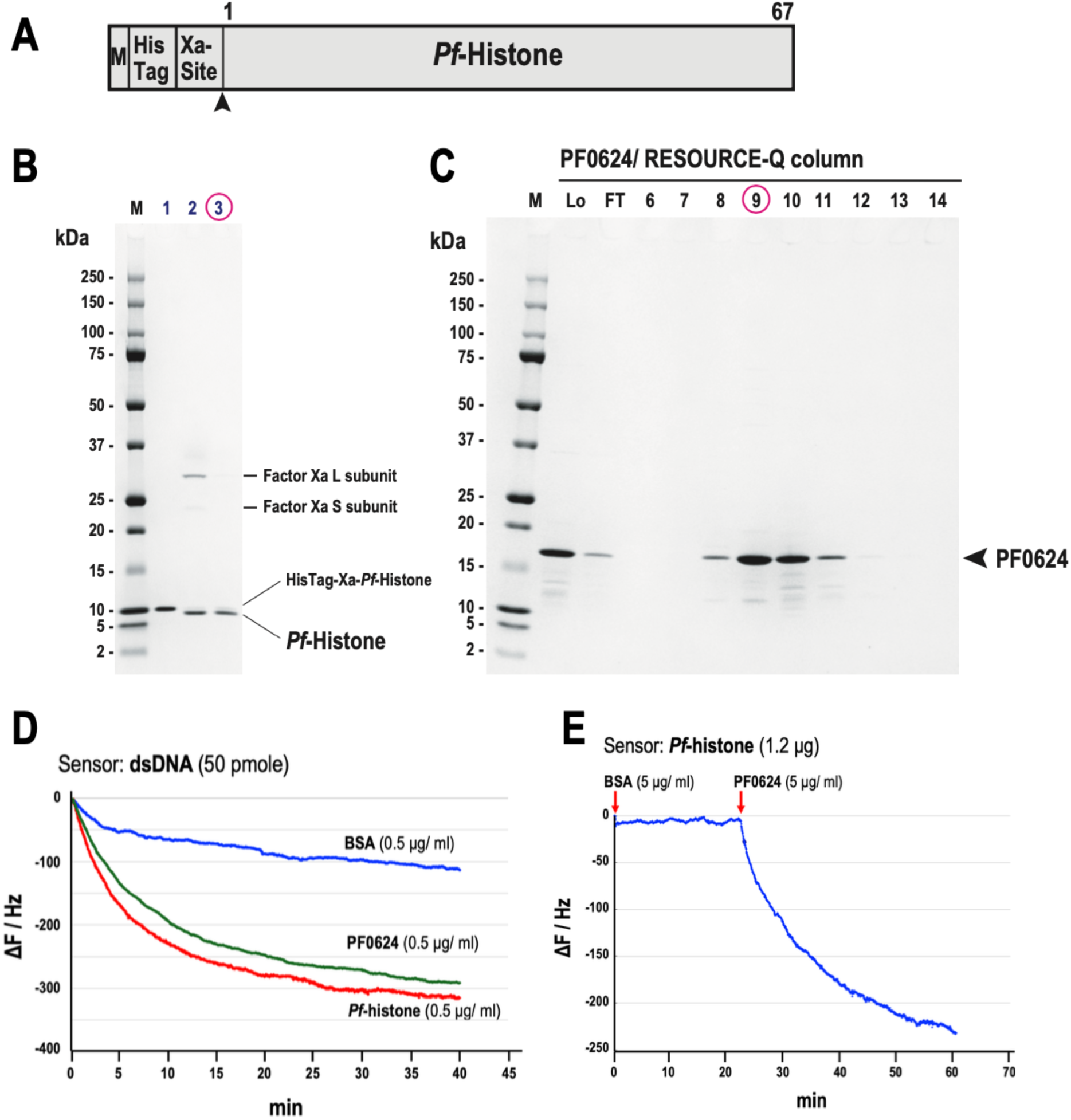
Analysis of the dsDNA-binding and *Pf*-histone-binding activities of PF0624 protein using the AffinixQ system. (A) Schematic of the recombinant *Pf*-histone protein used for the production and purification of the protein without a His-tag. M: methionine residue; His-tag: hexahistidine tag; Xa-site: target site for factor Xa. The numbers above the box indicate the positions of the amino acid residues. (B) Purification of *Pf*-histone without a His-tag. SDS-PAGE (10%‒20% polyacrylamide gels with Coomassie brilliant blue [CBB] staining) analysis of each purification step. Lane 1: fraction of purified His-tagged *Pf*-histone; Lane 2: fraction of His-tagged *Pf*-histone digested with Xa enzyme; Lane 3: fraction of purified *Pf*-histone. (C) Purification of PF0624 without a His-tag using RESOURCE-Q column chromatography. SDS-PAGE (10%‒20% polyacrylamide gels with CBB staining) analysis of each fraction from the column is shown. Lo: load fraction; FT: flow-through fraction. The arrowhead indicates the position of the purified protein. (D) Binding analysis of dsDNA and PF0624 protein using the AffinixQ system. (E) Binding analysis of *Pf*-histone and PF0624 protein using the AffinixQ system.

To investigate the DNA-binding properties of PF0624 under physiological conditions similar to the intracellular environment of *P. furiosus*, we conducted gel shift assays using a previously described method (33). Because greater protein amounts are required for this assay, we used His-tagged recombinant proteins. The addition of the His-tag did not alter the binding behavior observed in previous assays. The recombinant His-tagged PF0624 and *Pf*-histone proteins were expressed in *E. coli* and purified to near homogeneity, as confirmed by SDS-PAGE (Fig. 6A). Under DNA-binding conditions of 0.2 M KCl at 70°C, *Pf*-histone exhibited a clear dose-dependent shift of dsDNA, indicating its capacity to form archaeal chromatin-like structures (Fig. 6B). By contrast, no such shift was observed for BSA, which served as a negative control. Surprisingly, PF0624 alone did not induce a shift in dsDNA under the same conditions. However, when co-incubated with *Pf*-histone, PF0624 increased chromatin formation in a concentration-dependent manner (Fig. 6C,D). Moreover, this increase was temperature-dependent (Fig. 6E). At room temperature, the addition of 200 ng PF0624 increased chromatin formation by approximately 1.5-fold compared with the reactions lacking PF0624. By contrast, at 70°C, the same amount of PF0624 increased chromatin formation by more than threefold (Fig. 6F). This suggests that PF0624 is more effective in promoting chromatin assembly under high-temperature conditions, consistent with the hyperthermophilic nature of *P. furiosus*. The promotion of chromatin formation by PF0624 in the presence of *Pf*-histone did not require ATP, indicating that the effect of PF0624 does not depend on an energy-driven remodeling mechanism (Fig. 6G). Collectively, these findings suggest that PF0624 may act as a heat-activated chromatin assembly factor, cooperating with *Pf*-histone to facilitate nucleoid organization under extreme thermal stress.

**FIG 6.**
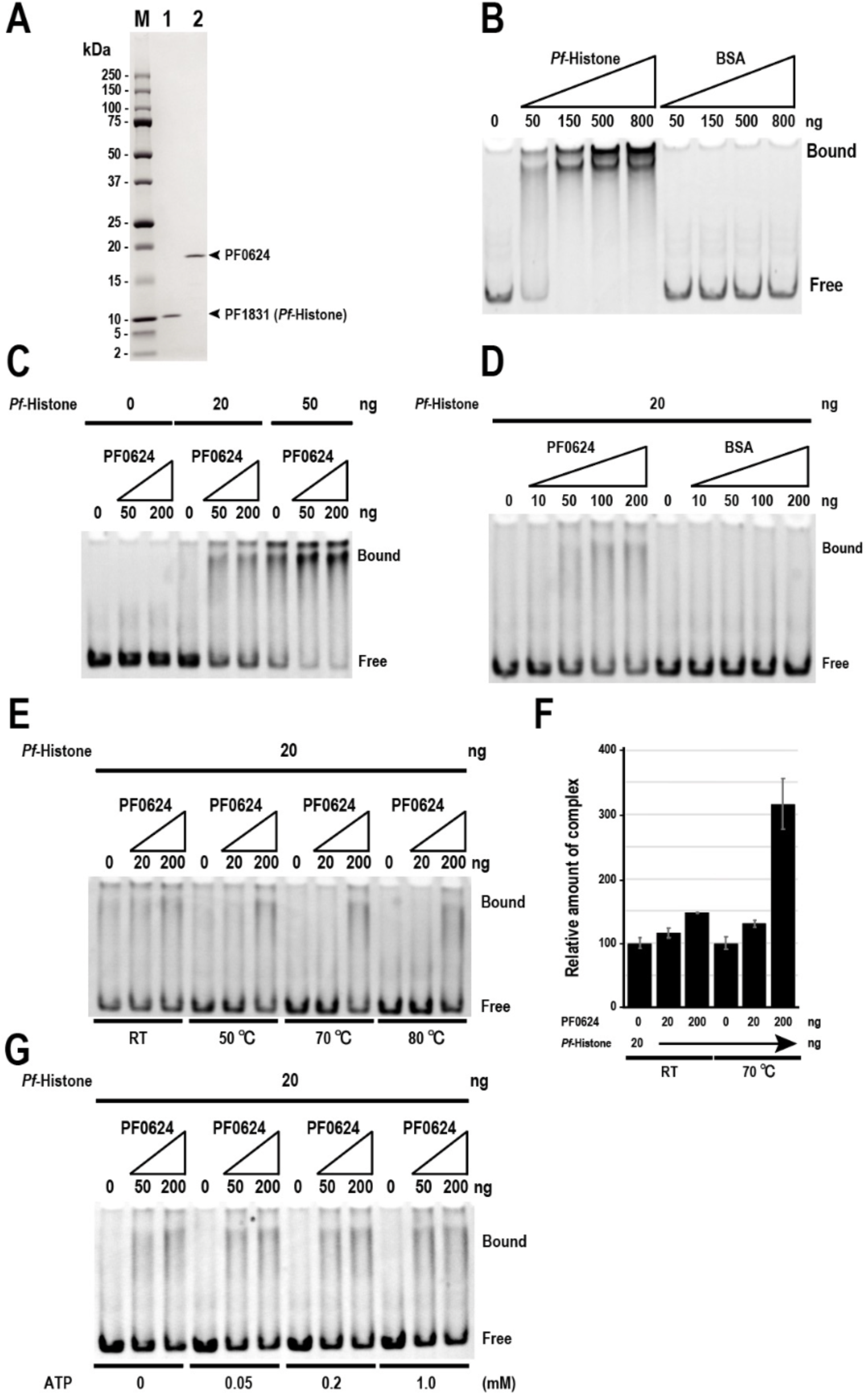
PF0624 enhances chromatin formation via *Pf*-histone. (A) Purification of recombinant His-tagged PF0624 and *Pf*-histone proteins. A peak sample from HiTrap chelating column chromatography, described in the Materials and Methods, was dialyzed and analyzed by SDS-PAGE with 10%‒20% polyacrylamide gels. The gel was stained with Coomassie brilliant blue. Arrowheads indicate the positions of each purified protein. M, molecular size markers (Bio-Rad). (B) Chromatin formation by *Pf*-histone. The gel shift assay for *Pf*-histone is shown. (C) PF0624 protein promotes chromatin formation by *Pf*-histone in a dose-dependent manner. (D) BSA does not affect chromatin formation by *Pf*-histone. (E) Effect of temperature on chromatin formation by *Pf*-histone via PF0624 protein. (F) Effect of PF0624 on promoting chromatin formation by *Pf*-histone is increased at higher temperatures. (G) ATP is not required for the effect of PF0624 protein on promoting chromatin formation by *Pf*-histone.

### Conclusions: Proposed model of the intracellular regulatory mechanisms in *P. furiosus* under heat shock stress

Based on our findings, we propose a model summarizing the regulatory events occurring in *P. furiosus* exposed to acute heat shock conditions. Even in hyperthermophilic archaea, acute heat shock triggers a series of rapid cellular responses within 15–45 min. The translation system is rapidly suppressed, as evidenced by a reduction in total tRNA, halving of the levels of 16*S* and 23*S* rRNAs and inhibiting precursor tRNA splicing. Despite this translational shutdown, approximately 90% of the proteins remain remarkably stable, even under boiling temperatures. One key exception is the cell division protein, FtsZ2, which undergoes a sharp, specific reduction in abundance. This suggests that cell division is halted during heat shock, allowing *P. furiosus* to endure high temperatures in a static cellular state. Concurrently, select genes are transcriptionally upregulated and their products efficiently translated, likely in anticipation of the damage caused by heat shock. These proteins include PF1883, a small heat shock protein functioning as a molecular chaperone; PF1385, a type-4 uracil-DNA glycosylase involved in DNA repair; and PF1616, a *myo*-inositol-1-phosphate synthase responsible for synthesizing di-*myo*-inositol-1-phosphate, a thermoprotectant. We also revealed a potential role for PF0624, a previously uncharacterized protein, in promoting *Pf*-histone-mediated chromatin assembly under high-temperature conditions. This activity may contribute to the stabilization of DNA and prevention of double-strand breaks during thermal stress. Recent findings have also linked histone-like proteins in *Halobacterium salinarum* to salt stress responses (34). Considering the diversity of histone-like proteins across archaea (31, 35, 36), further studies are warranted to clarify their broader roles in adaptation to environmental stress. At the transcriptional level, although many genes are upregulated in an apparently disordered fashion, the concomitant inhibition of translation prevents these increases in mRNA from being reflected at the protein level. Additionally, approximately 5% of transcripts are downregulated following heat shock, and half of those transcripts are associated with metabolic functions. This selective downregulation likely contributes to a cellular strategy to conserve resources and prioritize the expression and translation of stress-response genes during extreme heat exposure. Finally, the mechanisms underlying selective protein degradation and targeted translation of specific mRNAs under heat shock conditions remain largely unknown and require further investigation.

## MATERIALS AND METHODS

### P. furiosus culture

*P. furiosus* JCM 8422 (DSM 3638) was obtained from the RIKEN BioResource Center (Tsukuba, Japan) and cultured for 17–24 h at 95°C in 5 L of artificial seawater (Yashima Pure Chemicals, Osaka, Japan) containing KH_2_PO_4_ (2.5 g), yeast extract (5 g), Bacto tryptone (25 g), NiCl_2_·6H_2_O (10 mg), resazurin (5 mg), 50 mM Na_2_WO_4_·2H_2_O (1 mL), trace elements (50 mL), sulfur powder (2.5 g), and Na_2_S·9H_2_O (2.5 g) (final pH 6.5). The trace elements (pH 7.0) were 1.5 g/L of nitrilotriacetic acid, 3.0 g/L of MgSO_4_·7H_2_O, 0.64 g/L of MnSO_4_·5H_2_O, 1.0 g/L of NaCl, 0.1 g/L of FeSO_4_·7H_2_O, 0.18 g/L of CoSO_4_·7H_2_O, 76 mg/L of CaCl_2_, 0.18 g/L of ZnSO_4_·7H_2_O, 0.01 g/L of CuSO_4_·5H_2_O, 0.02 g/L of AlK(SO_4_)_2_·12H_2_O, 0.01 g/L of H_3_BO_3_, 0.01 g/L of Na_2_MoO_4_·2H_2_O, 0.025 g/L of NiCl_2_.6H_2_O, and 0.2 mg/L of Na_2_SeO_3_. The cells were harvested by centrifugation (6200 × *g* for 15 min at 4°C) and washed with buffer W containing 50 mM Tris-HCl (pH 8.0) and 10% (w/v) sucrose. The cell pellet was stored at −80°C. To induce heat shock, *P. furious* was cultured overnight at 90°C and then diluted 1:2 into 250 mL of fresh medium, and further cultured for 4 h at 90°C. The cells were boiled for 15, 30, and 45 min (corresponding to 101‒102°C at 1 atm) or not boiled (0 min, at 90°C), harvested by centrifugation (6200 × *g*, 5 min, 4°C), and washed with buffer W. The cell pellet was stored at −80°C until analysis.

### Total RNA preparation, northern blot analysis, and RT-quantitative PCR analysis

Total RNA was extracted from the *P. furiosus* cells using the RNeasy Midi Kit (Qiagen, Valencia, CA), with slight modification. Instead of using the RNeasy Midi column, we performed phenol– chloroform extraction to collect the complete set of RNAs efficiently. For northern blotting, total RNA (10 μg per lane) was separated on a denaturing 10% polyacrylamide gel containing 8 M urea and transferred onto a Hybond-N+ membrane (GE Healthcare, Piscataway, NJ) by electroblotting. The membranes were hybridized with the fluorescein amidite (FAM)-labeled synthetic DNA probe, 5tRNATrp40A-FAM for tRNA^Trp^(CCA) in PerfectHyb hybridization solution (Toyobo, Osaka, Japan) at 42°C overnight. After two washes each in 2× SSC (1× SSC: 0.15 M NaCl and 0.015 M sodium citrate) with 0.1% SDS for 5 min at room temperature and 0.2× SSC with 0.1% SDS for 10 min at 42°C, images were captured using a fluorescence imager (Molecular Imager FX Pro, Bio-Rad Laboratories, Hercules, CA). To analyze the mRNAs and rRNAs by RT-quantitative PCR (qPCR), RT was performed using the TaKaRa PrimeScript RT-PCR Kit (TaKaRa Bio Inc., Shiga, Japan), according to the manufacturer’s protocol. qPCR analysis was conducted using a real-time PCR System (MyGo Mini, IT-IS Life Science Ltd., Dublin, Republic of Ireland) with PerfeCTa SYBR Green FastMix (Quantabio, Beverly, MA), according to the manufacturers’ protocols. The relative amounts of mRNAs were normalized to the amount of 16*S* rRNA transcript in each sample.

### Proteomic analysis

The protein mixture was extracted by sonication (7 min) from pelleted *P. furiosus* with 100 mM triethylammonium bicarbonate (pH 8.5) containing 12 mM sodium deoxycholate, 12 mM sodium *N*-dodecanoyl sarcosinate, and 1% protease inhibitor cocktail (#P8340, Sigma-Aldrich, St. Louis, MO). The extract was centrifuged (18,000 × *g*, 10 min, 4°C) to remove any debris and dialyzed against phosphate-buffered saline (PBS)(-). Dialyzed proteins (55 μg) were incubated with benzonase (100 ng, MilliporeSigma, Darmstadt, Germany) for 5 min at 4°C. The proteins were then precipitated using trichloroacetic acid. Precipitated proteins were redissolved in guanidine hydrochloride, reduced with Tris(2-carboxyethyl)phosphine, alkylated with iodoacetamide, and digested with lysyl endopeptidase and trypsin. The digested peptides were cleaned up using a C18 monospin-column (GL Science Inc., Tokyo, Japan). Then, the purified peptides (500 ng) were analyzed using a nano liquid chromatography system (Easy-nLC 1200, Thermo Fisher Scientific, Waltham, MA) coupled to a mass spectrometer (Q-Exactive HF-X, Thermo Fisher Scientific). Peptides were separated on a C18 separation column (Nikkyo Technos, Tokyo, Japan; 3 μm, 100 µm inner diameter × 12 cm). Mobile phases A and B consisted of 0.1% formic acid in water and 0.1% formic in 80% acetonitrile, respectively. We used an 80-min gradient at a constant flow rate of 300 nL/min, ranging from 5% to 40% mobile phase B. The eluent was directly introduced into the mass spectrometer, which was operated in data-dependent acquisition mode to select the top 25 precursor ions. Mass spectra were recorded from 380 to 1500 *m/z*. Survey scans were acquired at a resolution of 60,000 at 200 *m/z*, and the resolution for tandem mass spectra was set to 15,000 at 200 *m/z*. All data were analyzed using Proteome Discoverer 2.2 (Thermo Fisher Scientific) with the SEQUEST search engine. All mass spectra were searched against protein sequences within the RefSeq *P. furiosus* DSM 3638 protein database (NCBI). The false discovery rate was set at 1% for the peptide‒spectrum match. All quantitations performed in this study were done at the peptide level. Label-free quantification was performed based on precursor signal intensity.

### Transcriptome analysis

Before performing RNA-seq analysis, archaeal 16*S* and 23*S* rRNAs were removed from the total RNA using the Ribominus transcriptome isolation kit (Thermo Fisher Scientific). Because this kit is not specialized for archaeal rRNA removal, we designed and chemically synthesized 5ʹ-biotinated oligonucleotides, Biotin-16*S*80-A and Biotin-23*S*80-A (Eurofins, Luxembourg, Luxembourg), were used as antisense oligonucleotides for the 16*S* and 23*S* rRNAs of *P. furiosus*. RNA-seq was conducted by GeneBay Inc. (Yokohama, Japan). Sequencing was performed on a sequencing system (NovaSeq 6000, Illumina, San Diego, CA) using 2 × 150 paired-end sequence reads. The control sample yielded 11,965,139 reads. The samples obtained after 15, 30, and 45 min of heat shock yielded 12,018,192 reads, 11,090,151 reads, and 12,632,403 reads, respectively. *P. furiosus* contains 2127 coding DNA sequences according to the NCBI GenBank file (RefSeq ID: NC_003413.1) (37). After removing the adapter and low-quality sequences from the transcriptome sequencing data, genome mapping and quantification of the expression levels were performed. Adapter and low-quality sequence removal was conducted in paired-end mode with Trimmomatic-0.33 Mapping (38). Expression levels were quantified using bowtie2-2.0.0.0-beta7 (39) and RSEM 1.3.1 (40) with the gff3 file (https://bacteria.ensembl.org/Pyrococcus_furiosus_dsm_3638_gca_000007305/Info/Index). Overall, 11,746,477 reads for the control sample, and 11,899,498 reads, 10,710,786 reads, and 12,394,290 reads for samples obtained after 15, 30, and 45 min of heat shock, respectively, were mapped onto the genome. Of the generated data, we used the data contained in the transcripts per million file in the subsequent analysis.

### Estimation of gene function using arCOG

The functional estimation and classification of *P. furiosus* genes and proteins was performed using arCOG. arCOG is a database of the function of archaeal genes and proteins, which are divided into 25 functional classifications according to the COG class (41). Genes and proteins that are not assigned to any COG class are classified as “Poorly Characterized.” Supplementary Tables S1 and S2 list the functional classifications of the genes and proteins examined in this study.

### Prediction of the three-dimensional (3D) structure of the PF0624 protein

Using the amino acid sequence of the PF0624 protein (GenBank ID: AAL80748.1) derived from *P. furiosus* strain DSM 3638, we performed homology modeling of the protein using Phyre2 (Protein Homology/analogY Recognition Engine V 2.0) in normal mode (29). UCSF Chimera version 1.15 was used to draw the 3D structure (42). Because the crystal structure of archaeal histone B derived from *M. fervidus* (PDB ID: 5T5K) has been published (30), we used it for comparison with the PF0624 protein structure. The MatchMaker command in UCSF Chimera was used with the default settings to superimpose the two structures.

### Construction of expression vectors

Expression vectors to produce recombinant His-tagged PF0624 (a hypothetical or histone-fold protein) and His-tagged PF1831 (*Pf*-histone type A) in *E. coli* were constructed as described previously (17). Genomic DNA from *P. furiosus* JCM 8422 (DSM 3638) was isolated using a GNOME DNA Isolation Kit (MP Biomedicals, Ohio, USA) and partially digested with Sau3AI. The resulting DNA fragments were separated by electrophoresis on a 0.7% (w/v) agarose gel, and fragments of approximately 5–15 kb were extracted and used as PCR templates. Each gene was amplified by PCR and cloned into the pET-23b expression vector (Novagen, Madison, WI) using site-specific primers. These primers were designed to introduce *Nhe*I and *Xho*I sites for the PF0624 gene, and *Nde*I and *Xho*I sites for the PF1831 gene at the 5′ and 3′ termini, respectively. The resulting plasmids encoded each protein with a C-terminal His-tag. The inserted sequences were verified by DNA sequencing and matched the corresponding entries in the NCBI database for *PF0624* (accession no.: AAL80748) and *PF1831* (accession no.: AAL81955).

To generate a His-tag-free version of PF0624, a stop codon was introduced into the 3′ N-terminal antisense primer to prevent translation of the His-tag sequence present in the pET-23b vector. For PF1831, the N-terminal His-tag was cleaved post purification using factor Xa protease (Novagen) (see Fig. 5A). The corresponding coding sequence was artificially synthesized (Eurofins Genomics) with codon optimization for *E. coli*. The synthetic gene included *Nde*I and *Xho*I restriction sites at its 5′ and 3′ ends, respectively, and was subcloned into the same sites in pET-23b. The structures of the artificially synthesized nucleotide sequences and the corresponding predicted protein products are shown in Supplementary Fig. S3. The oligoribonucleotides used in this study are listed in Supplementary Table S3.

### Expression and purification of recombinant proteins

The His-tagged recombinant PF0624 and PF1831 proteins were prepared essentially as described previously (19). Briefly, *E. coli* BL21(DE3) cells were transformed using the respective expression plasmids. The transformants were grown at 37°C in Luria–Bertani medium supplemented with 50 µg/mL ampicillin until the mid-log phase and were then induced with 0.4 mM isopropyl-β-D-thiogalactoside (IPTG). After 14–16 h of incubation at 30°C, the cells were harvested by centrifugation (9000 × *g*, 15 min, 4°C) and lysed by sonication (3–4 min) in Proteus binding buffer (50 mM sodium phosphate, pH 7.4, 300 mM NaCl, 10 mM imidazole). The lysate was heat-treated at 85°C for 15 min to denature endogenous *E. coli* proteins, followed by centrifugation (18,000 × *g*, 10 min, 4°C) to remove debris. The supernatant was applied to a HiTrap chelating column (GE Healthcare, Chicago, IL), and the eluted recombinant proteins were dialyzed against buffer D (50 mM Tris-HCl, pH 8.0, 1 mM EDTA, 0.02% [v/v] Tween 20, 7 mM 2-mercaptoethanol, and 10% [v/v] glycerol).

The His-tag-free PF0624 protein, which was highly expressed in *E. coli*, was purified by heat treatment and anion exchange chromatography. After IPTG induction, the cells were harvested and total proteins were extracted using a buffer containing 50 mM Tris-HCl (pH 8.0), 1 mM EDTA, 0.02% (v/v) Tween 20, 7 mM 2-mercaptoethanol, 10% (v/v) glycerol, and 1 M NaCl. The extract was heat-treated at 85°C for 15 min, centrifuged (18,000 × *g*, 10 min, 4°C), and dialyzed against buffer D. The dialyzed lysate was applied to a 1 mL RESOURCE-Q column (Cytiva, Marlborough, MA) equilibrated with buffer D and eluted with a linear NaCl gradient (0–1.0 M) using an ÄKTA Purifier FPLC system (GE Healthcare). The elution peak was detected in fractions corresponding to 300–400 mM NaCl, and fraction #9 was used in this study (Fig. 5C). To prepare His-tag-free *Pf*-histone, the precursor protein with an N-terminal His-tag was first extracted and heat-treated, similar to the procedure described above. The elution peak was collected using a HiTrap chelating column and dialyzed against PBS (-). Factor Xa cleavage and purification were then performed using the Factor Xa Cleavage Capture Kit (Merck Millipore, Burlington, MA) according to the manufacturer’s protocol.

### Molecular interaction analysis using the biosensor system

The binding interactions among purified PF0624 protein, *Pf*-histone, and dsDNA were analyzed using a biosensor system (AffinixQ, Piezo Parts Co., Ltd., Tokyo, Japan), which employs the quartz crystal microbalance method to detect molecular binding on a quartz sensor with nanogram-level sensitivity by measuring mass changes. To analyze the interaction between dsDNA and the proteins, 50 pmol of thiolated dsDNA (see Supplementary Table S3) was immobilized on the sensor surface to enhance binding. The reaction was carried out at 45°C in binding buffer A containing 10 mM Tris-HCl (pH 8.0), 0.5 mM EDTA, 2.5 mM MgCl_2_, and 150 mM NaCl, supplemented with PF0624 protein or *Pf*-histone at a final concentration of 0.5 μg/mL. The change in frequency (Δ*F* [Hz]) of the quartz crystal resonator was monitored for approximately 40 min, until it reached equilibrium. For the *Pf*-histone–PF0624 interaction, *Pf*-histone (1.2 μg) was first immobilized on the sensor. Then, BSA was then added to binding buffer A to a concentration of 5 μg/mL to verify minimal nonspecific interaction at 45°C. After confirmation, PF0624 was added and Δ*F* was monitored until the binding had stabilized.

### Gel shift assay of the archaeal histone‒DNA complex and effect of PF0624 protein on chromatin formation

To form the archaeal histone‒DNA complexes, binding reactions containing the PCR-amplified dsDNA (part of *P. furiosus* genomic region, 462 bp, 100 ng) and purified PF1831 protein (*Pf-* histone, 0‒800 ng) were pre-incubated at room temperature for 10 min in 20 µL of gel-shift binding buffer (10 mM Tris/HCl [pH 7.5], 50 mM NaCl, 200 mM KCl, 0.5 mM EDTA, 0.5 mM MgCl_2_, 5% [v/v] glycerol, and 1 mM dithiothreitol). To examine the effect of PF0624 protein, the purified PF0624 protein (0‒200 ng) was added to the archaeal histone–DNA complex solutions, and the mixtures were incubated at 70°C for 10 min. The reaction was stopped by adding 1/10 vol of 10× blue dextran dye (4 mM Tris/HCl [pH 7.5], 0.4 mM EDTA, 20% [v/v] glycerol, and a small amount of blue dextran). The DNA–histone complexes were promptly separated by electrophoresis on a 6% (w/v) nondenaturing polyacryl-amide gel at room temperature and visualized with ethidium bromide staining. The DNA–histone complexes were quantified by scanning the fluorescent image using an FX Pro computerized image analyzer (Bio-Rad Laboratories).

## ACKNOWLEDGEMENTS

The authors thank all the members of the RNA Group at the Institute for Advanced Biosciences of Keio University, Japan for their insightful discussions.

## AUTHORS’ CONTRIBUTIONS

Haruko Okabe, Conceptualization, Data curation, Formal analysis, Investigation, Methodology, Validation, Visualization, Writing – original draft | Masahiro C. Miura, Conceptualization, Data curation, Formal analysis, Investigation, Methodology, Validation, Visualization, Writing – original draft | Asako Sato, Formal analysis, Investigation, Validation | Shungo Adachi, Formal analysis, Investigation, Writing – original draft | Akio Kanai, Conceptualization, Formal analysis, Funding acquisition, Investigation, Supervision, Validation, Writing – original draft, Writing – review and editing.

## DATA AVAILABILITY

The raw RNA sequences have been deposited in the DDBJ database and are available under BioProject accession no. PRJDB13268 (Run ID: DRR355548–DRR355551). The raw proteomic mass spectrometry data have been deposited in the Japan ProteOme STandard Repository (jPOST) under the accession no. JPST003726.

## SUPPLEMENTAL MATERIAL

Supplemental material is available for this article: Supplementary Tables S1–S3 and Supplementary Figures S1–S3.

